# Genomic analysis of *Megalocytivirus* genomes reveals widespread recombination

**DOI:** 10.1101/2025.10.27.684589

**Authors:** Polly I. Hannaford, Lachlan Coff, Robyn N. Hall, Jeffrey Go, Nicholas J. G. Moody, Robert Lanfear

## Abstract

Megalocytiviruses are pathogens of global significance that can lead to substantial economic losses in aquaculture. Recombination among megalocytiviruses is typically assumed to be rare, although it has been relatively understudied. Here, we uncover widespread recombination within megalocytiviruses through detailed analyses of 63 *Megalocytivirus* genomes, including two which are newly sequenced and assembled. We also identify a number of genes which megalocytiviruses have likely obtained from outside the family *Iridoviridae* (iridovirids). These results have serious implications for the biosecurity management of megalocytiviruses, as they indicate that *Megalocytivirus* strains could be misclassified based on traditional approaches which target individual loci in the genome. We use this new knowledge of recombination to estimate updated phylogenetic trees of megalocytiviruses at the family-, genus-, and species-level. These trees show strong support for the designation of two novel species within the genus *Megalocytivirus* and highlight the difficulty of placing highly recombinant genomes in a single phylogenetic framework. We discuss the implications of our work for disease management, and the importance of genome-wide recombination detection and phylogenomic analysis in the classification and genetic characterisation of megalocytiviruses.

## Introduction

Megalocytiviruses are members of the genus *Megalocytivirus* (family *Iridoviridae*) which are among the most significant pathogens affecting finfish aquaculture worldwide. Infection with these double-stranded DNA viruses can lead to 100% mortality in a broad range of finfish of commercial and cultural importance (Kurita et al. 2002; Kurita and Nakajima 2012; Kawato et al. 2017). The host and geographical range of megalocytiviruses is rapidly expanding with outbreaks in Asia, Africa, the United States, South America, and Europe (Go et al. 2006; Lopez-Porras et al. 2018; Halaly et al. 2019; Figueiredo et al. 2022; Alathari et al. 2024). The World Organisation for Animal Health (WOAH) is the international standard-setting body for trade in animal commodities, and lists one species of *Megalocytivirus*, *Megalocytivirus pagrus1*, as a pathogen requiring global management to minimise the risk of transboundary spread (World Organisation for Animal Health 2025).

The phylogeny of megalocytiviruses is well characterised. Most studies rely on the analysis of the major capsid protein (MCP) gene or the adenosine triphosphatase (ATPase) gene (Do et al. 2004; Lü et al. 2005; Shi et al. 2010; Nolan et al. 2015; Kim et al. 2019; Senapin et al. 2019; Sun et al. 2025). Such analyses suggest that there are two species that are part of the genus *Megalocytivirus*: *M. pagrus1* and *M. lates1*. *Megalocytivirus pagrus1* genomes are typically classified as one of three genotypes (or ‘genogroups’) – red seabream iridovirus (RSIV), infectious spleen and kidney necrosis virus (ISKNV), or turbot reddish body iridovirus (TRBIV) (Kawato et al. 2020). These genotypes typically share more than 95% nucleotide sequence similarity and are often further split into two ‘clades’ or ‘subtypes’ (Koda et al. 2018; Kawato et al. 2020; Fusianto et al. 2023; Kushala et al. 2024). *Megalocytivirus lates1* encompasses a single genotype known as scale drop disease virus (SDDV) (Fu et al. 2021; ICTV 2024). Threespine stickleback iridovirus (TSIV) (Waltzek et al. 2012; Yoxsimer et al. 2024) and European chub iridovirus (ECIV) (Halaly et al. 2019) have been proposed as novel *Megalocytivirus* species (Waltzek et al. 2012; Halaly et al. 2019; Yoxsimer et al. 2024). Phylogenomic analyses have largely confirmed the phylogeny from single-gene studies (de Groof et al. 2015; Koda et al. 2018; Koda et al. 2019; Jeong et al. 2021; Fusianto et al. 2023; Liao et al. 2023). Although, such multi-gene studies can only include the limited number of samples for which complete (or near-complete) genome sequences are available. Further, no previous study has incorporated genomes from *M. pagrus1, M lates1*, TSIV, and ECIV, meaning we lack a phylogenomic perspective on the *Megalocytivirus* genus as a whole (de Groof et al. 2015; Koda et al. 2018; Halaly et al. 2019; Koda et al. 2019; Liao et al. 2023; Yoxsimer et al. 2024).

Despite extensive research on the phylogeny of megalocytiviruses, recombination among *Megalocytivirus* genomes is yet to be fully investigated and would have serious implications for classification and disease risk management. Phylogenetically derived taxonomic designations are used to manage the risk of megalocytiviruses (WOAH 2024). For example, mandatory disease reporting to animal health authorities and treatment strategies can be specific to the species or genotype (Kahn et al. 1999; Dong et al. 2017; World Organisation for Animal Health 2025) and clade designations based on the analysis of a single loci have been used to infer the suspected pathway of introduction of megalocytiviruses (Tsai et al. 2005; Nolan et al. 2015; Girisha et al. 2020; Zhu et al. 2021; Azad et al. 2024). Recombination can occur when a host cell is co-infected with two or more pathogens, and genetic information is swapped between them (Pérez-Losada et al. 2015). Alternatively, genetic information can be shared directly between host and viral genomes (Kurita and Nakajima 2012). If recombination is common in megalocytiviruses, relying on individual loci to classify virus genotypes could be misleading. Additionally, the concatenation of recombinants in multi-gene phylogenies can distort evolutionary signals (Gori et al. 2016). Finally, the recommended approaches for confirming *Megalocytivirus* genotypes largely rely on the PCR amplification of a single gene (Crane and Moody 2018; World Organisation for Animal Health 2025). If DNA sequences targeted by PCR assays are involved in recombination events, this could undermine standard approaches for identifying megalocytiviruses and responding to disease outbreaks (Wen et al. 2013; Dohál et al. 2021; Tu et al. 2024).

There have been isolated reports of recombination in *Megalocytivirus* genomes, although the overall extent of recombination in this group remains unclear. RSIV-Ku, isolated in Penghu Taiwan from cage-cultured red seabream (*Pagrus major*), was classified as ISKNV based on the analysis of the ORF037L gene (Shiu et al. 2018). Later analysis revealed a section of the genome shares a greater level of similarity to RSIV (Kawato et al. 2020). Additionally, two genomes isolated from moribund farmed barramundi (*Lates calcarifer*) in Thailand were found to have a ’reticular pattern’ with ISKNV and RSIV genomes (Kerddee et al. 2021). The MCP gene of African lamprey iridovirus, which is classified as *M. pagrus1*, shares a greater level of homology with ISKNV whilst the ATPase gene is more similar to RSIV, indicative of recombination (Sudthongkong et al. 2002; Fonseca Jr et al. 2023). Mixed RSIV sub-type I/II strains 17SbTy and SB5-TY have been isolated from farmed fish in Korea, both classified based on the analysis of PCR amplicons (Kim et al. 2019; Jeong et al. 2021). The existence of such putatively recombinant genomes has led to a call for new classifications to resolve phylogenetic inconsistencies (Kang et al. 2024). Recombination has also been documented in the family *Iridoviridae* (Kurita and Nakajima 2012; Vilaça et al. 2019; Li et al. 2024). Notably, megalocytiviruses appear to have inherited the small subunit of ribonucleotide reductase (RR-2) gene from a past eukaryotic host, whilst the remaining iridovirids are thought to have acquired theirs from rickettsia-like eubacteria (Kurita and Nakajima 2012).

In this study, we used a comprehensive catalogue of existing and new *Megalocytivirus* genomes to examine evidence of recombination. We used a whole genome approach to test for recombination between the genotypes and clades of *M. pagrus1*, and phylogenetic methods to test for recombination between *Megalocytivirus* species, and between megalocytiviruses and other iridovirids and more distantly related taxa. Our results show that recombination between clades and genotypes of *M. pagrus1* is widespread, with recombination occurring throughout the genome. In addition, we provide new evidence that megalocytiviruses have acquired genetic material from non-viral species. Finally, we discuss the implications of these findings for our understanding of the evolutionary history and biosecurity management of megalocytiviruses.

## Results

### Widespread evidence of recombination among *Megalocytivirus pagrus1* genotypes and sub-clades

Using a variety of recombination detection methods and highly conservative criteria for determining recombination, we found compelling evidence of widespread recombination among *M. pagrus1* genomes. We annotated a dataset of 58 high quality genomes from *M. pagrus1* (see Materials and Methods and supplementary table 1) and used RDP4 (version 101) (Martin et al. 2015) to test for signals of recombination in a whole genome alignment (see Materials and Methods and Supplementary Materials). We detected recombination in 14 of the 58 genomes (24%; see Figure 1). Of these, nine genomes (16%) showed evidence of recombination between different genotypes, and ten (17%) showed evidence of recombination between clades within a genotype (Figure 1). Placing these events in a phylogenetic context suggests that there were at least six independent recombination events between genotypes (Figure 1). The recombinant genomes were isolated from farmed food fish (except for three recombinant genomes, for which the source was unknown; Figure 1, supplementary table 1). This was despite the inclusion of 17 genomes isolated from ornamental fish (Figure 1, supplementary table 1). Further details on each recombination event are provided on GitHub at this link: https://github.com/PollyHannah/Phylogenomic-study/blob/main/recombination/recombination_results_refined.xlsx.

**Figure 1.**
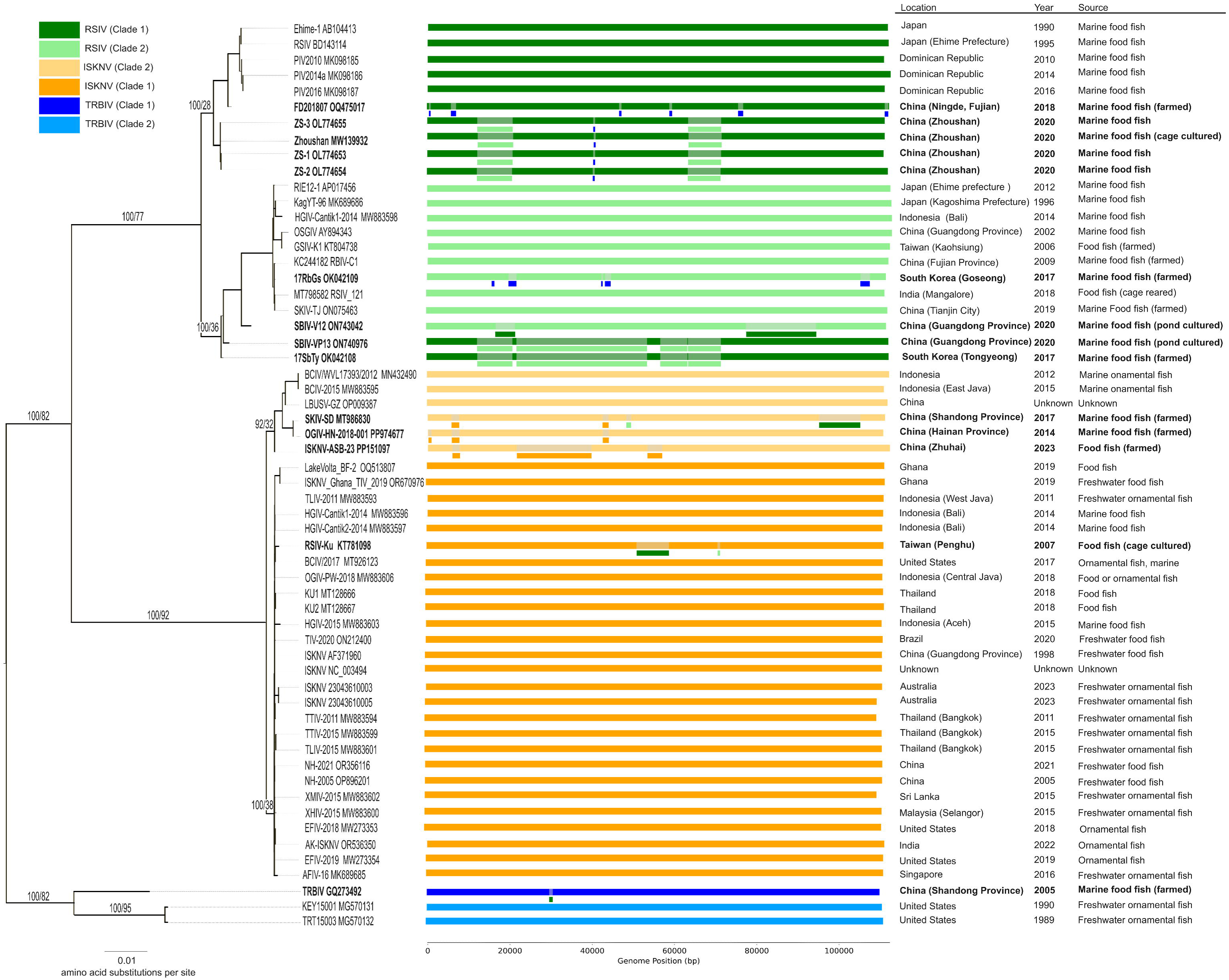
Phylogenetic tree of genomes from *Megalocytivirus pagrus1*. This maximum likelihood tree is based on 68 concatenated genes. Branch leaves are annotated with isolate name and GenBank accession number. Branch labels show bootstrap values followed by gene concordance factors for key branches. Coloured bars represent genomes coloured according to clade designations based on the major capsid protein (MCP) gene. The shaded sections in the bars represent putative recombinant regions within genomes, with the acquired sequences represented by short bars beneath the shaded sections (coloured according to clade designations based on the MCP gene). The location, year, and source (type of fish host) are provided for each genome. The data for putatively recombinant genomes is bolded.

### No evidence of recombination between *Megalocytivirus* species

The degree of divergence between *Megalocytivirus* species precluded the generation of a whole-genome alignment. We therefore relied on phylogenetic approaches for detecting recombination between species. To do this, we added five genomes to the dataset – three from *M. lates1*, one from ECIV and one from TSIV – resulting in 63 genomes. We used OrthoFinder (version 2.5.4) (Emms and Kelly 2015; Minh et al. 2020) to extract orthologous loci that were present in at least 70% of genomes and built phylogenetic trees in IQ-TREE (version 2.2.0.5) (Minh et al. 2020) (see Materials and Methods). This resulted in phylogenetic trees for 116 loci. Detailed examination of each single-locus tree showed no evidence of strongly supported topological differences that could be the result of recombination (supplementary table 2). In a handful of trees, the sole TSIV sequence was placed within the *M. pagrus1* clade, but in all cases the TSIV sequence was on a long terminal branch, and the placement was close to the root of the *M. pagrus1* clade, consistent with phylogenetic misplacement of divergent sequences and unlikely to be the result of intra-species recombination. All genus-level gene trees are provided on GitHub at this link https://github.com/PollyHannah/Phylogenomic-study/tree/main/iqtree_genus_trees.

### Evidence of recombination between megalocytiviruses and other genera

We used the same phylogenomic approach to test for evidence of recombination between *Megalocytivirus* genomes and other iridovirid species or more distantly related taxa. First, we supplemented the 63-genome dataset with ten representative genomes from the six other genera from the family *Iridoviridae* (*Ranavirus*, *Lymphocystivirus*, *Iridovirus*, *Chloriridovirus*, *Decapodiridovirus* and *Daphniairidovirus*). We then applied the same filtering, alignment, orthology detection, and phylogenetic tree building steps as above, resulting in 27 locus trees (see ‘Materials and Methods’). We manually examined each gene tree for evidence of recombination and found compelling evidence of recombination in three loci; RR-2, transcription factor S, and kinase (supplementary table 2). In all of these loci the *Megalocytivirus* clade was connected to the remaining iridovirid genomes by a surprisingly long branch. The mean branch length connecting the *Megalocytivirus* clade to the other iridovirid genera across all 27 loci was 0.8 amino acid substitutions per site, but it was more than double that in each of these three loci. To examine this further, we performed BLAST analyses of the genes in our dataset. For most genes in our dataset, the BLAST analysis showed that the *Megalocytivirus* gene is most closely related to sequences from other iridovirids, as expected. However, for the three genes identified as connecting to the iridovirids on long branches in the phylogenetic analyses, the BLAST analysis revealed that the *Megalocytivirus* sequences were more similar to sequences from distantly related taxa. The *Megalocytivirus* RR-2 gene is most similar to sequences from eukaryotic organisms including fish. The *Megalocytivirus* transcription factor S gene is most similar to sequences from eukaryotic organisms including protists and fungi. The *Megalocytivirus* kinase gene is most similar to sequences from bacteria. Details of these BLAST results are provided in the Supplementary Materials.

### A new phylogenomic framework for *Megalocytivirus*

Finally, we estimated maximum likelihood phylogenomic trees for megalocytiviruses. We estimated separate trees for each taxonomic level (species, genus, and family). At each taxonomic level, we excluded loci with evidence of recombination since these could mislead estimates of the relationships among genomes. At the species level, we excluded all loci with evidence of recombination between genotypes. We were unable to exclude loci with evidence of recombination between clades, as this would have excluded the majority of loci. Phylogenomic analysis of 68 orthologues from *M. pagrus1* resulted in a tree with strong support and high concordance for all three *M. pagrus1* genotypes (all bootstrap values 100%; gene concordance factors 82% for TRBIV, 92% for ISKNV, and 77% for RSIV), and high bootstrap but lower concordance factors for the two sub-clades of each genotype (Figure 1). At the family level, a phylogenomic analysis of 24 non-recombinant loci recovered the *Megalocytivirus* genus as a sister clade to *Lymphocystivrus* and *Ranavirus* with 100% bootstrap support and a gene concordance factor of 95% (Figure 2A). At the genus level, a phylogenomic analysis of 96 non-recombinant loci showed 100% bootstrap support and high concordance factors for all major groupings, including for the grouping of TSIV with *M. pagrus1*, and the grouping of ECIV with *M. lates1* (Figure 2B).

**Figure 2.**
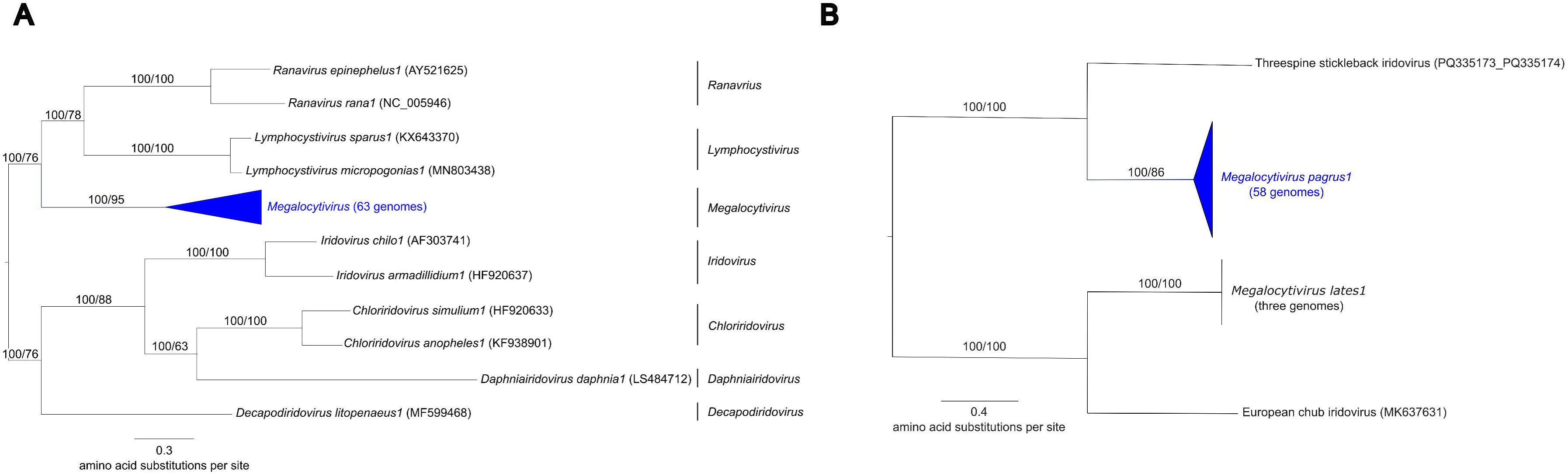
Concatenated maximum likelihood phylogenetic trees for (A) the family *Iridoviridae* and (B) the genus *Megalocytivirus*. Branch leaves are annotated with isolate name and GenBank accession number. Branch labels show bootstrap values followed by gene concordance factors. The family level tree (A) is based on 24 concatenated genes and is rooted according to the International Committee for the Taxonomy of Viruses Virus Taxonomy Profile for *Iridoviridae* (Chinchar et al. 2017). The genus-level tree (B) is based on 96 concatenated genes and is rooted according to the family level tree (A). The slightly lower gene concordance factor for *Megalocytivirus pagrus1* results from a handful of loci in which a threespine stickleback iridovirus gene is placed within the *Megalocytivirus pagrus1* clade (see main text).

## Discussion

This study reveals widespread recombination among *Megalocytivirus* genomes. Phylogenetic and whole-genome recombination detection methods revealed recombination within and between all three *M. pagrus1* genotypes (Figure 1, supplementary table 2). It is documented that fish can be simultaneously infected with multiple sub-clades of *Megalocytivirus* genotypes (Fu et al. 2023), providing the necessary conditions for intra-genotypic recombination. We also identified three genes that megalocytiviruses appear to have obtained from distantly related non-viral species (supplementary table 2, Supplementary Materials). The RR-2 gene of megalocytiviruses has previously been shown by Kurita and Nakajima (2012) to share little homology to that of all other iridovirids. Their analysis suggests that the *Megalocytivirus* genus acquired the RR-2 gene from eukaryotes, a result consistent with our finding that this gene shows closer similarity to eukaryotic homologs. Further, our analysis reveals that the *Megalocytivirus* transcription factor S and kinase genes are also likely of eukaryotic and bacterial origin respectively. These findings suggest that *Megalocytivirus* genomes have a previously underappreciated proclivity to acquire genetic material from non-viral species.

Our findings significantly advance our understanding of the extent and nature of recombination among *Megalocytivirus* genomes. Previous evidence of recombination among megalocytiviruses was limited to RSIV-Ku, a natural recombinant of ISKNV and RSIV (Shiu et al. 2018; Kawato et al. 2020) and sporadic reports of ‘mixed type’ genomes (Sudthongkong et al. 2002; Kim et al. 2019; Jeong et al. 2021; Kerddee et al. 2021). Our analysis reproduced the finding of recombination in RSIV-Ku, and confirmed the presence of recombination in ‘mixed type’ strain 17SbTy, for which a whole genome was available (Jeong et al. 2021) (Figure 1). We provide strong evidence for six independent inter-genotypic recombination events across the three genotypes of *M. pagrus1* (Figure 1). This included putative recombination between RSIV and TRBIV genomes collected from South Korea and China, where these pathogens circulate in the same species (Shi et al. 2004; Shi et al. 2010; Kim et al. 2019; Fusianto et al. 2023). Likewise, ISKNV and RSIV cause disease in the same species of fish in China and Taiwan, making the existence of recombinant strains plausible (Wen and Hong 2016; Zhu et al. 2021; Fu et al. 2023; Fusianto et al. 2023; Choi et al. 2024). Intriguingly, all putatively recombinant genomes whose origin is known were isolated from food fish (mainly farmed marine fish) despite the inclusion of a large number of genomes from freshwater ornamental fish (Figure 1, supplementary table 1). Marine farming practices like cage culture expose fish to wild animals (Kawato et al. 2021; Kawato et al. 2023) and multiple species are sometimes cultured in the same location at high densities. This makes cross-species transmission and co-infection more likely, which may facilitate the emergence of genetic recombinants (Do et al. 2005; Wang et al. 2022; Cai et al. 2024; Liu et al. 2025).

Our results show that only analysing a small number of genes can lead to misclassification of megalocytiviruses. Classification of megalocytiviruses often relies on the sequencing the MCP gene and/or the ATPase gene (Kurita et al. 2002; Shi et al. 2010; Kawato et al. 2020; Fu et al. 2021). The ATPase gene appears to have been swapped between genomes in different *M. pagrus1* clades making it an unreliable target for the clade-level classification of this species (supplementary table 2). Whilst our findings confirm the MCP as one of the most reliable loci for classifying megalocytiviruses (supplementary table 2), classification based on a single locus ignores the presence of large recombinant regions. This is problematic not only because it misrepresents the composition of the genome, but also because recombination can influence virulence (Jeong et al. 2021). For instance, Jeong et al. (2021) reported a much lower survival rate in rock bream (*Oplegnathus fasciatus*) infected with the RSIV-TRBIV recombinant 17RbGs compared with the RSIV mixed sub-type 17SbTy. Crucially, in the recombinant 17RbGs isolate, and in many of our samples, the MCP and

ATPase gene sequences used to classify megalocytiviruses provide no evidence that the genotype is recombinant. Full genome sequencing, however, reveals that the genome is recombinant and can explain differences in virulence and other viral characteristics (Jeong et al. 2021). The presence of highly recombinant genomes also challenges the traditional clade designations used for *M pagrus1.* For example, strains such as SBIV-VP13 and 17SbTy have genomes that are mosaics of clade one and clade two genotypes of RSIV (Figure 1). This not only makes a clade designation questionable, but results in an intermediate phylogenomic placement between clade one and clade two, as shown in Figure 1. Together, our findings highlight an urgent need for routine whole genome sequencing and analysis for the accurate classification and genetic characterisation of *Megalocytivirus* strains.

Recombination appears to have minimal impact on phylogenomic estimates derived from multi-gene datasets. We sought to estimate phylogenies that avoided, as far as is possible, the inclusion of loci that were involved in recombination. This resulted in estimating phylogenetic trees from different sets of loci than previous studies (supplementary table 2). Despite this, our phylogenies are highly congruent with previously-published phylogenetic trees for megalocytiviruses (de Groof et al. 2015; Koda et al. 2018; Koda et al. 2019; Jeong et al. 2021; Koda et al. 2021; Zhao et al. 2022; Fusianto et al. 2023; Liao et al. 2023). Although, multi-gene phylogenies showing the relationships between *M. pagrus1* genotypes are most often presented as cladograms (Koda et al. 2018; Koda et al. 2021; Fusianto et al. 2023). Whilst the topology of the species-level tree generated in this study reflects that of those previously published, the use of cladograms precludes comparisons of the evolutionary distances between taxa (Brower 2016). At the family-level, recombination appears to have little effect on multi-gene phylogenies (de Groof et al. 2015; Jeong et al. 2021; Zhao et al. 2022; Liao et al. 2023); this is despite putatively recombinant genes being included in the 26 ‘core genes’ identified by Eaton et al. (2007), widely used to phylogenetically analyse iridovirid taxa (supplementary table 2) (de Groof et al. 2015; Koda et al. 2018; Jeong et al. 2021; Liao et al. 2023). It is possible that the conflicting phylogenetic signals caused by recombinant loci are ‘drowned out’ by the collective signal of the remaining data, as has been observed in previous studies (Tonini et al. 2015; Gori et al. 2016). Importantly, the family-level tree provides strong support for the designation of *Daphniairidovirus* as a novel iridovirid genus (Figure 2A), which is not currently recognised by the International Committee on Taxonomy of Viruses (ICTV). Similarly, the genus-level tree provides strong support the designation of two novel *Megalocytivirus* species to accommodate ECIV and TSIV (Figure 2B).

Recombination in megalocytiviruses has serious implications for disease management, potentially affecting classification, cell line susceptibility, vaccine efficacy and pathogen virulence. Most molecular diagnostic assays for megalocytiviruses target sequences within the MCP gene (Rimmer et al. 2012; Kawato et al. 2021; Moody et al. 2022). Although some target other regions such as the putative myristoylated protein (Lin et al. 2017), for which evidence of recombination was shown in this study (supplementary table 2). The analysis of PCR amplicons from such recombinant regions could lead to the misidentification of megalocytiviruses (Fusianto et al. 2023). It is possible that recombination could be partially responsible for the observed lack of cross-protection provided by *Megalocytivirus* vaccines, where recombinant genes are targeted in their development (Dong et al. 2017; Fu et al. 2023). Further, the observed inter-genotypic recombination among strains may explain the strain-specific patterns of cell line susceptibility observed for *M. pagrus1* (Kwon et al. 2011; Wen et al. 2013). Finally, the rapid evolution of viruses through recombination may be an important risk factor where it leads to increased levels of virulence and pathogenicity or the emergence of novel viruses with potentially altered disease phenotypes (Jeong et al. 2021). Overall, this work highlights the need for continuous monitoring of genomes to document and track new genetic variants. This is not only to anticipate shifts in virus phenotype, but also to accurately trace recombinant strains sharing high levels of sequence similarity to divergent taxa.

In summary, our study provides strong evidence of recombination in conserved genes within the genus *Megalocytivirus* and beyond. We detected recombination and re-estimated the phylogenetic relationships among *Megalocytivirus* taxa and other members of the family *Iridoviridae*. The information provided will guide future research on the evolutionary history of megalocytiviruses, and the biosecurity management of these pathogens.

## Materials and Methods

### Data collection and review

We first collected available public data on completely sequenced *Megalocytivirus* and other iridovirid genomes from the National Centre for Biotechnology Information (NCBI) database. A total of 79 genomes were collected and reviewed, including two novel genomes (see Novel genomes). All sequences deposited as whole genomes under the genus *Megalocytivirus*, and ten representative genomes from the six other *Iridoviridae* genera (*Ranavirus*, *Lymphocystivirus*, *Iridovirus*, *Chloriridovirus*, *Decapodiridovirus* and *Daphniairidovirus*) were collected from the NCBI database. The genomes from genera other than *Megalocytivirus* were chosen to span the deepest node of the clade as in Zhao et al. (2022).

Species currently recognised by the ICTV as part of the genus *Megalocytivirus* are *M. pagrus1* and *M. lates1*. Two additional putative *Megalocytivirus* genomes, ECIV and TSIV, were included in this study. Neither are currently classified by the ICTV. The TSIV genome was present as two separate contigs (accessions PQ335173 and PQ335174) which we concatenated in the same order as was done in Yoxsimer et al. (2024), henceforth referred to as TSIV (PQ335173_PQ335174).

All *Megalocytivirus* genomes were classified to the genotype level based on sequence similarity using FastANI (version 1.34) (Jain et al. 2018) which computes the average nucleotide identity (ANI) between query genomes and reference sequences. For *Megalocytivirus* reference sequences we used: ISKNV (ISKNV, accession AF371960), RSIV (KagYT-96, accession MK689686), TRBIV (TRBIV, accession GQ273492), SDDV (SDDV-ZH-06/20, accession OM037668), TSIV (TSIV, accession PQ335173_PQ335174), and ECIV (ECIV, accession MK637631). The ANI of each genome is available on GitHub (https://github.com/PollyHannah/Phylogenomic-study/blob/main/taxonomy.csv).

Genomes were excluded from the final dataset if there was evidence of sequencing or assembly errors or having too many stop codons in annotations (see the Annotation section). An RSIV genome (accession KC138898) and *Lymphocystis disease virus 1* genome (accession L63545) were excluded due to not being the expected length. The latter genome was replaced in the dataset with a *Lymphocystis disease virus 3* genome (accession KX643370).

### Novel genomes

#### Virus isolation method

Two novel megalocytiviruses were isolated from ornamental fish, swordtail (*Xiphophorus helleri*) and southern platyfish (*X. maculatus*), which screened positive by quantitative PCR (qPCR) for megalocytiviruses in November 2023. Tissue homogenate samples of ISKNV-positive tissue from each fish were submitted to the CSIRO Australian Centre for Disease preparedness (ACDP) Fish Disease Laboratory for disease investigations.

Cell cultures of the SKF-9 cell line (Kawato et al. 2017) were used in this study at passage numbers 36 to 38. The cell cultures were maintained at 25 °C in Hank’s minimum essential medium (Thermo Fisher Scientific) and monitored for cytopathic effect. The tissue culture supernatant was harvested from each sample and tested by qPCR to confirm virus isolation results.

#### Genome sequencing and assembly

Samples of extracted DNA (5 μl) from the virus isolated from *X. helleri* and *X. maculatus* were used to generate short-read sequencing libraries using the Nextera® XT DNA Library Preparation Kit and Index Kit v2 Set A (Illumina®). The libraries were sequenced using a P1 600-cycle cartridge on a NextSeq™ 2000 with onboard basecalling using BCL Convert (version 4.0.3). Separate sequencing libraries were constructed for each sample. The quality of the raw paired-end reads was checked using FastQC (version 0.12.1) (Brown et al. 2017). Reads of low quality were trimmed using Fastp (version 0.23.2) (Chen 2023) with reads with a base quality lower than 20 or a length less than 100 bp discarded. The reads from the cell line host (*O. punctatus)* used to grow the virus were removed using Bowtie (version 2.5.4) (Langmead and Salzberg 2012) and SAMtools (version 1.18.0) (Danecek et al. 2021) using the reference genome for *O. punctatus* (accession CNP0001488) obtained from the China National GeneBank DataBase https://db.cngb.org/.

The unmapped reads generated from the samples sourced from *X. helleri* and *X. maculatus* were partitioned into four and two groups respectively due to the large size of the datasets. *De novo* assembly of the paired end reads for each group was done using SPAdes (version 3.15.5) (Prjibelski et al. 2020) and Unicycler (version 0.5.1) (Wick et al. 2017). Contigs of the expected size were retained and contigs sharing high sequence similarity to host genomes were discarded. To confirm they were identical, the retained contigs were aligned in Geneious Prime (version 2020.5.2) using the progressiveMauve algorithm (version 1.1.3) (Darling et al. 2010) with default parameters.

The two novel genomes from the virus isolates obtained from *X. helleri* and *X. maculatus* were named 23-04361-0003 and 23-04361-0005 respectively. A total of 23 M reads were generated for 23-04361-0003 and 25.6 M reads for 23-04361-0005, with 2.7% of reads trimmed due to low quality for both sets of reads. The reads for 23-04361-0003 assembled into one contig 111,083 bp in length which shared 99.88% identity to ISKNV genome with the accession AF371960. The reads for 23-04361-0005 assembled into two contigs. One of the two contigs shared 99.6% identity to the mitochondrial genome of the host cell line. The remaining contig 110,927 bp in length was accepted as the genome given it shared 99.92% identity to the ISKNV genome with the accession AF371960. The reads have been deposited to the GenBank Sequence Read Archive (SRA) under accessions SRR35394460 for 23-04361-0003, and SRR35415016 for 23-04361-0005. The genome sequences for 23-04361-0003 and 23-04361-0005 have been deposited in GenBank under the accession numbers PX411171 and PX411172 respectively.

### Annotation

To ensure consistent annotation across all samples, all genomes used in this study were annotated with Prokka (version 14.5) using default parameters with the kingdom set to viruses (Seemann 2014). All annotations were then manually checked against previously assigned annotations (where available) from the NCBI database. We followed different procedures for *Megalocytivirus* genomes versus those from other genera.

To annotate *Megalocytivirus* genomes, we chose a single reference genome from each genotype which was published in a peer reviewed journal and was fully annotated on Genbank – these are the same genomes as in the ‘Data Collection and review’ section above. This was done to generate a manually annotated reference genome from each genotype, with which to annotate the remaining genomes. For each of these reference genomes, we compared the annotations assigned by Prokka against the corresponding annotations in the GenBank file for that genome. Where the two annotations matched, we kept that annotation. When the annotations differed, we did a BLASTp (Altschul et al. 1990) search on each annotation option and kept the annotation with a higher number of matches, percent identity to matches, and percent query cover. The only exception was if the annotation that scored best in the BLASTp analysis had > 5 stop codons while the alternative had fewer, in which case we kept the lesser scoring option. Where there were no matches for a Prokka annotation and no gene present at or near that interval in the annotated GenBank genome, we removed the Prokka annotation. We only added GenBank genome annotations which were not identified by Prokka, if a BLASTp search returned at least one match for the Genbank annotation. The same process was completed for all *Megalocytivirus* genomes entered into GenBank as unclassified at the genotype level, of which there were four (accessions MG570131, MG570132, OQ475017, and OL310752).

Each of the remaining *Megalocytivirus* genomes were annotated by circularising the genomes and then transferring annotations from a database containing each of the manually annotated reference genomes using the ‘Annotate From’ function in the ‘Live Annotate and Predict’ menu in Geneious Prime (version 2020.2.5). To do this, the percent similarity was set at 85%, meaning only annotations sharing a minimum of 85% similarity with the reference sequence were transferred.

For genomes in genera other than the genus *Megalocytivirus*, a sub-set of 25 Prokka-assigned annotations were manually checked to avoid manually checking all loci. The subset of genes chosen were those present in all genomes as identified by running OrthoFinder (version 2.5.4) with standard options (Emms and Kelly 2019). The chosen annotations were checked and edited using the same decision-making criteria and process described above.

Annotations were then manually curated. Annotations for putative genes not starting with the amino acid methionine were discarded. Where there were ≤ 5 stop codons in an annotation, the stop codons were removed from the annotation. Where there were > 5 stop codons in an annotation, the annotation was removed. Where there were > 5 annotations removed from a genome, the genome was excluded from the analysis due to potential sequencing or assembly error(s).

A total of 108 updates to annotations were made across the reference and unclassified genomes. Four genomes were excluded due to assumed sequencing or assembly errors following re-annotation. Red sea bream iridovirus genomes (RBIV-KOR-TY1, accession AY532606), large yellow croaker iridovirus (accession AY779031), and SDDV genome (C4575, accession NC_027778) were excluded due to there being > 5 annotations in the genome removed. Unclassified genome Zhoushan, from the *Megalocytivirus* genus (accession OL310752), was excluded due to a highly conserved gene being split into two genes in disparate locations in the genome.

The number of Open reading frames (ORFs) identified in each genome included in the study is provided in supplementary table 1. Between 115 and 122 ORFs were found in *M. pagrus1* genomes with a minimum of 132 identified in *M. lates1* genomes. A total of 114 and 103 ORFS were found in ECIV and TSIV genomes respectively and between 97 and 278 in genomes from all other genera. All re-annotated sequences can be found on GitHub at this link: https://github.com/PollyHannah/Phylogenomic-study/blob/main/annotated_genome_sequences.geneious.

### Final dataset

The final set of genomes included in the study comprised 73 genomes. This included 58 genomes from *M. pagrus1*, three from *M. lates1*, one each from TSIV and ECIV, and ten from iridovirid genera other than the genus *Megalocytivirus* (supplementary table 1). Only three *M. pagrus1* genomes of the TRBIV genotype were available for inclusion in this study.

### Orthologue detection

Next, loci were identified which could be used for phylogenetic analyses. We used OrthoFinder (version 2.5.4) (Emms and Kelly 2019) to identity sets of orthologous genes (orthogroups) at the family-level (all 73 iridovirid genomes), genus-level (all 63 megalocytivirus genomes) and species-level (all 58 *M. pagrus1* genomes). We ran OrthoFinder (version 2.5.4) (Emms and Kelly 2019) on each taxonomic level dataset to generate multiple sequence alignments (MSAs) using MAFFT (version 7.49) (Katoh and Standley 2013) which included genes from 70% of the genomes in each dataset. We chose an ‘occupancy threshold’ of ≥ 70% to maximise the number of loci with potentially useful genetic information. We identified a total of 473, 187 and 129 orthogroups using OrthoFinder at the family-, genus- and species-levels. Of these, 16, 49 and 82 were present in 100% of genomes at the family-, genus- and species-level respectively. These numbers increased to 115, 116 and 116 when we reduced the occupancy threshold to ≥ 70%. Of the orthogroups we identified at the family level, 84 no longer contained sequence information unique to the genus-level alignments and were therefore discarded (we discarded a further four family-level orthogroups following the alignment editing step, see below). Of the orthogroups identified at the genus level, 20 did not contain sequence information unique to the species-level alignments and were therefore discarded. This resulted in 31, 96 and 116 genes being retained for analysis at the family-, genus- and species-levels.

We realigned the MSAs for every locus using Muscle5 (version 5.0.1428) (Edgar 2004), and manually edited each locus to improve alignments and remove poorly aligned sequences. Where two sequences from the same taxon were allocated to an orthogroup, we only retained the sequence which returned the highest percent identity and the largest number of sequence matches from a BLASTp search (Altschul et al. 1990). We removed a total of 25, seven and four sequences from the family-, genus- and species-level alignments respectively, and made 20 manual edits to the family-level alignments. Following this, an additional four family-level alignments no longer contained sequence information unique to the genus-level alignments, and we therefore discarded them, leading to 27 genes being retained at the family level. The manual edits to MSA files can be found on GitHub at this link: https://github.com/PollyHannah/Phylogenomic-study/blob/main/alignment_manual_changes.xlsx.

We trimmed the manually edited MSAs to remove columns with less than 25% occupancy using TrimAL (version 1.4.1r22) (Capella-Gutiérrez et al. 2009). We determined the possible identity of genes in each orthogroup by using the BLAST+ suite (version 2.16.0) (Camacho et al. 2009) to run a BLASTp search (Altschul et al. 1990), comparing the genes against proteins in the Swiss-Prot database and selecting those with the highest Score (Bits) and the lowest E-value. Where the putative identities were unlikely, we revised them based on the results of a BLASTp search (Altschul et al. 1990). The putative identity of these genes can be found in supplementary table 2.

### Gene tree estimation

To generate gene trees, we used QMaker in IQ-TREE (version 2.4.0) (Minh et al. 2021) to estimate amino acid substitution models which best fit the data. We built one model from the genus level MSAs (Q.mcv) and another from the family level MSAs (Q.iridoviridae). We then generated a maximum likelihood gene tree for each alignment using IQ-TREE 2 (version 2.2.0.5) (Minh et al. 2020) with 1000 ultrafast bootstrap replicates using ultrafast bootstrap approximation (Hoang et al. 2018). We used ModelFinder (Kalyaanamoorthy et al. 2017) in IQ-TREE (version 2.2.0.5) (Minh et al. 2020) to determine the best-fit substitution model for each analysis from the following list of 30 potential models: Blosum62, cpREV, Dayhoff, DCMut, FLAVI, FLU, HIVb, HIVw, JTT, JTTDCMut, LG, mtART, mtMAM, mtREV, mtZOA, mtMet, mtVer, mtInv, PMB, Q.bird, Q.insect, Q.mammal, Q.pfam, Q.plant, Q.yeast, rtREV, VT, WAG, Q.iridoviridae, Q.mcv.

### Recombination analysis

We used whole-genome recombination detection methods to test for recombination among *M. pagrus1* genotypes and clades. Investigating recombination among *M. lates1* genomes was deemed unnecessary given the species is represented by a single genotype and clade. To identify putative inter- and intra-genotypic recombination in *M. pagrus1* genomes using RDP4 (version 101) (Martin et al. 2015), we generated a whole genome alignment of *M. pagrus1* nucleotide sequences using the progressiveMauve plugin (version 1.1.3) (Darling et al. 2010) in Geneious Prime (version 2020.5.2) with default parameters. The five recombination detection methods used within RDP4 were RDP (Martin et al. 2015), GENECONV (Padidam et al. 1999), MAXCHI (Smith 1992), CHIMAREA (Posada and Crandall 2001) and 3Seq (Boni et al. 2007). The sequences were specified as linear and the option ‘Disentangle overlapping events’ was unselected, as recommended for datasets with > 50 sequences (Martin 2021). All other RDP4 parameters were set to default.

We took a conservative approach to refining preliminary recombination hypotheses identified by RDP4 (version 101) (Martin et al. 2015). We only accepted events if a) they were detected by the five detection methods used, b) the recombination signal was visible in the plots generated using the MAXCHI (Smith 1992) and CHIMAREA (Posada and Crandall 2001) (example provided in Supplementary Materials), and c) the recombination could be observed by comparing the topology of two phylogenetic trees – one based on the putative recombinant region and the other on combined regions where recombination was not detected (example provided in Supplementary Materials), and d) the recombination could be clearly identified in the whole genome alignment. These criteria were based on recommendations in the RDP4 Instruction Manual (Martin 2021).

We did not employ whole genome recombination detection methods at the family or genus levels given whole genome alignments were not feasible due to the level of divergence between genomes. Instead, we used phylogenomic approaches to identify loci which may have been involved in recombination. To do this, we analysed each individual locus tree at the family- and genus-level looking for two telltale signs of recombination. First, we looked for strong phylogenetic support for non-monophyly of the *Megalocytivirus* genus in the family-level gene trees, or non-monophyly of any *Megalocytivirus* species in the genus-level trees. Where we found potential evidence of non-monophyly we performed an AU test (Shimodaira 2002) in IQ-TREE (version 2.2.0.5) (Minh et al. 2020) to determine whether monophyly could be statistically rejected. No evidence of recombination was reported for loci where monophyly could not be statistically rejected. Second, we looked for evidence of surprisingly long internal branches which connected the *Megalocytivirus* genus to other iridovirid genera in the family-level gene trees, or connecting *Megalocytivirus* species in the genus-level gene trees. These long branches can occur if the *Megalocytivirus* genomes had acquired sequences from distantly related (e.g. non-viral) taxa (Kurita and Nakajima 2012). When we observed a long branch, we used tBLASTp (Altschul et al. 1990) to compare a *Megalocytivirus* sequence from that locus to all sequences contained in NCBI GenBank database. We did this to identify close relatives to the *Megalocytivirus* sequences which were not included in our alignments. No evidence of recombination was reported for loci where a tBLASTp (Altschul et al. 1990) search returned no hits for taxa outside of the genus *Megalocytivirus*.

### Phylogenetic analysis

We used IQ-TREE (version 2.2.0.5) (Minh et al. 2020) to build a phylogenomic tree at the species-, genus-, and family-level. For the family- and genus-level analyses, we only used genes that were not identified as putatively recombinant. For the species-level analyses, genes showing evidence of inter-genotypic recombination (recombination between genotypes) were excluded. However, genes showing evidence of intra-genotypic recombination (recombination within *M. pagrus1* clades) were included. We included genes showing intra-genotypic recombination at species-level to maximise the volume of genomic sequence information used to estimate genetic relationships between genotypes.

For all three analyses, we built a concatenated maximum likelihood tree with 1000 bootstraps using ultrafast bootstrap approximation (Hoang et al. 2018), and used ModelFinder (Kalyaanamoorthy et al. 2017) to determine the best-fit substitution model (as above). We then re-estimated gene trees for select loci (as above) and used IQ-TREE (version 2.2.0.5) (Minh et al. 2020) to calculate gene concordance factors for every branch in the concatenated tree. All trees were annotated and formatted for publication using FigTree (version 1.4.4) (Rambaut 2018) and Inkscape (version 1.4.2).

## Code Availability

The code, data, and step-by-step instructions required to replicate this analysis are provided on GitHub at this link: https://github.com/PollyHannah/Phylogenomic-study.

## Supporting information

Supplementary table 1_Genome summary table

Supplementary table 2_Gene summary table

Supplementary data_Kinase gene_BLAST+

Supplementary data_RR-2 gene_BLAST+

Supplementary data_Transcription factor S_BLAST+

## Acknowledgements

This study was funded by the CSIRO ACDP Infectious Animal Diseases and Zoonoses Program. Polly Hannaford was supported by the Australian Department of Agriculture Fisheries and Forestry through a Sir Roland Wilson Scholarship. The authors thank the Elizabeth Macarthur Agricultural Institute for the initial identification of ISKNV, and subsequent submission of the ISKNV-positive tissue samples, from which the novel genomes were sequenced. The authors thank AFDL for the virus isolation work, and Dr James O’Dwyer for his involvement in whole genome sequencing.

## Conflict of interest

All authors declare no competing interests.

## Notes

https://github.com/PollyHannah/Phylogenomic-study

### Competing Interest Statement

The authors have declared no competing interest.

