## Supplementary table 1_Genome summary table for "Genomic analysis of *Megalocytivirus* genomes reveals widespread recombination"

**Supplementary table 1.** Genomes included in this study. **No^o^** Number **MCP** Major Capsid Protein **bp** Base pairs **ORFs** Open Reading Frames assigned in this study **N/A** Not applicable **ISKNV** Infectious spleen and kidney necrosis virus **RSIV** Red sea bream iridovirus **TRBIV** Turbot reddish body iridovirus **SDDV** Scale drop disease virus **N/A** Not applicable **FV3** Frog virus 3 **LYCIV** Large yellow croaker iridovirus **SKIV** Spotted knifejaw iridovirus **PIV** pompano iridovirus **RBIV** Red bream iridovirus **AFIV** Angel fish iridovirus **GSIV** Giant seaperch iridovirus **OSGIV** Orange spotted groper iridovirus **LBUSV** Large-mouth bass ulcerative syndrome virus **TSGIV** Three spot gourami iridovirus **SGIV** Singapore groper iridovirus **TSIV** Three-spine stickleback iridovirus **DIV-1** *Daphnia iridescent virus 1*. **LCDV** Lymphocystis disease virus **IIV** Invertebrate iridescent virus **AMIV** Anopheles minimus iridovirus.

| Genus | Species | Genotype | Isolate or strain | Accession Number | Host | Host type | Source | Year collected | Location collected | Genome size (bp) | ORFs | References |
| --- | --- | --- | --- | --- | --- | --- | --- | --- | --- | --- | --- | --- |
| *Megalocytivirus* | *Megalocytivirus pagrus1* | ISKNV | ISKNV | AF371960 | Mandarin fish (*Siniperca chuatsi*) | Food fish, freshwater | Farm | 1998 | China, Nanhai, Guangdong Province | 111,362 | 121 | He et al. (2001) |
|  |  |  | AFIV-16 | MK689685 | Angelfish (*Pterophyllum scalare*) | Ornamental fish, freshwater | Ornamental fish trade | 2016 | Singapore | 111,127 | 122 | Kawato et al. (2020) |
|  |  |  | BCIV/WVL17393/2012 | MN432490 | Banggai cardinalfish  (*Pterapogon kauderni*) | Ornamental fish, marine | Unknown | 2012 | Indonesia | 112,618 | 118 | GenBank |
|  |  |  | BCIV/2017 | MT926123 | Banggai cardinalfish  (*Pterapogon kauderni*) | Ornamental fish, marine | Unknown | 2017 | United States | 111,920 | 122 | GenBank |
|  |  |  | SKIV-SD | MT986830 | Spotted knifejaw (*Oplegnathus punctatus)* | Food fish, marine | Farmed | 2017 | China, Shandong Province | 111,198 | 120 | Huang et al. (2021) |
|  |  |  | LBUSV-GZ | OP009387 | Unknown | Unknown | Unknown | Unknown | China | 112,248 | 121 | GenBank |
|  |  |  | ISKNV-ASB-23 | PP151097 | Barramundi (*Lates calcarifer*) | Food fish, marine or freshwater | Farmed | 2023 | China, Zhuhai | 112,236 | 120 | Sun et al. (2024) |
|  |  |  | TIV-2020 | ON212400 | Tilapia (*Oreochromis niloticus*) | Food fish, freshwater | Unknown | 2020 | Brazil | 111,837 | 121 | Alathari et al (2023) |
|  |  |  | ISKNV_Ghana_TIV_2019 | OR670976 | Tilapia (*Oreochromis niloticus*) | Food fish, freshwater | Unknown | 2019 | Ghana | 111,831 | 120 | GenBank |
|  |  |  | BCIV-2015 | MW883595 | Banggai cardinalfish (*Pterapogon kauderni*) | Ornamental fish, marine | Unknown | 2015 | Indonesia, Ketapang, East Java | 111,666 | 120 | Fusianto et al. (2023) |
|  |  |  | KU2 | MT128667 | Barramundi (*Lates calcarifer*) | Food fish, marine or freshwater | Unknown | 2018 | Thailand | 111,610 | 119 | GenBank |
|  |  |  | OGIV-PW-2018 | MW883606 | Giant gourami (*Osphronemus goramy*) | Food fish and ornamental fish, freshwater | Unknown | 2018 | Indonesia, Purwokerto,Central Java | 111,609 | 121 | Fusianto et al., (2023) |
|  |  |  | KU1 | MT128666 | Barramundi (*Lates calcarifer*) | Food fish, marine or freshwater | Unknown | 2018 | Thailand | 111,487 | 119 | GenBank |
|  |  |  | EFIV-2019 | MW273354 | Rainbow shark (*Epalzeorhynchos frenatum*) | Ornamental fish | Unknown | 2019 | United States | 111,380 | 121 | GenBank |
|  |  |  | EFIV-2018 | MW273353 | Rainbow shark (*Epalzeorhynchos frenatum*) | Ornamental fish | Unknown | 2018 | United States | 111,369 | 120 | GenBank |
|  |  |  | ISKNV | NC_003494 | Unknown | Unknown | Unknown | Unknown | Unknown | 111,362 | 122 | He et al. (2001) |
|  |  |  | LakeVolta_BF-2 | OQ513807 | Tilapia (*Oreochromis niloticus*) | Food fish | Unknown | 2019 | Ghana | 111,362 | 118 | Alathari et al. (2023) |
|  |  |  | TTIV  -2015 | MW883599 | Three spot gourami (*Trichopodus trichopterus*) | Ornamental fish, freshwater | Unknown | 2015 | Thailand, Bangkok | 111,351 | 122 | Fusianto et al., (2023) |
|  |  |  | NH-2005 | OP896201 | Mandarin fish (*Siniperca chuatsi*) | Food fish, freshwater | Unknown | 2005 | China | 111,315 | 121 | GenBank |
|  |  |  | NH-2021 | OR356116 | Mandarin fish (*Siniperca chuatsi*) | Food fish, freshwater | Unknown | 2021 | China | 111,315 | 121 | GenBank |
|  |  |  | TLIV-2011 | MW883593 | Pearl gourami (*Trichogaster leeri*) | Ornamental fish, freshwater | Unknown | 2011 | Indonesia, Bekasi, West Java | 111,309 | 121 | Fusianto et al., (2023) |
|  |  |  | RSIV-Ku | KT781098 | Red seabream (*Pagrus major*) | Food fish | Farmed, cage culture | 2007 | Taiwan, Penghu | 111,154 | 122 | Shiu et al. (2018) |
|  |  |  | HGIV-2015 | MW883603 | Hybrid groper (*Epinephelus fuscoguttatus x Epinephelus polyphekadion*) | Food fish, marine | Unknown | 2015 | Indonesia, Lhokseumawe, Aceh | 111,138 | 120 | Fusianto et al., (2023) |
|  |  |  | HGIV-Cantik1-2014 | MW883596 | Hybrid groper (*Epinephelus fuscoguttatus x Epinephelus polyphekadion*) | Food fish, marine | Unknown | 2014 | Indonesia, Bali | 111,039 | 120 | Fusianto et al., (2023) |
|  |  |  | HGIV-Cantik2-2014 | MW883597 | Hybrid groper (*Epinephelus fuscoguttatus x Epinephelus polyphekadion*) | Food fish, marine | Unknown | 2014 | Indonesia, Bali | 111,039 | 120 | Fusianto et al., (2023) |
|  |  |  | TLIV-2015 | MW883601 | Dwarf gourmani (*Trichogaster lalius*) | Ornamental fish, freshwater | Unknown | 2015 | Thailand, Bangkok | 111,014 | 120 | Fusianto et al., (2023) |
|  |  |  | XMIV-2015 | MW883602 | Southern platysfish (*Xiphophorus maculatus*) | Ornamental fish, freshwater | Unknown | 2015 | Sri Lanka | 110,962 | 119 | Fusianto et al., (2023) |
|  |  |  | XHIV-2015 | MW883600 | Green swordtail (*Xiphophorus hellerii*) | Ornamental fish, freshwater | Unknown | 2015 | Malaysia, Selangor | 110,894 | 120 | GenBank |
|  |  |  | OGIV-HN-2018-001 | PP974677 | Grouper (*Epinephelus spp.*) | Food fish, marine | Farmed | 2014 | China, Hainan Province | 110,699 | 118 | Cao et al. (2024) |
|  |  |  | AK-ISKNV | OR536350 | Angelfish (*Pterophyllum scalare*) | Ornamental fish | Unknown | 2022 | India | 110,460 | 117 | Kushala et al. (2025) |
|  |  |  | TTIV-2011 | MW883594 | Blue gourami (*Trichopodus trichopterus*) | Ornamental fish, freshwater | Unknown | 2011 | Thailand, Bangkok | 110,394 | 118 | Fusianto et al. (2023) |
|  |  |  | ISKNV | 23043610003 | Swordtail (*Xiphophorus helleri*) | Ornamental fish, freshwater | Imported ornamental fish | 2023 | Australia (imported fish) | 111,083 | 120 | This study |
|  |  |  | ISKNV | 23043610005 | Platys (*Xiphophorus maculatus*) | Ornamental fish, freshwater | Imported ornamental fish | 2023 | Australia (imported fish) | 110,927 | 118 | This study |
|  |  | RSIV | KagYT-96 | MK689686 | Japanese amberjack (*Seriola quinqueradiata*) | Food fish, marine | Unknown | 1996 | Japan, Kagoshima Prefecture | 112,710 | 120 | Kawato et al. (2020) Ito et al. (2013) |
|  |  |  | OSGIV | AY894343 | Orange spotted grouper  (*Epinephelus coioides*) | Food fish, marine | Farmed | 2002 | China, Guangdong Province | 112,636 | 117 | Lü et al., (2005) |
|  |  |  | RSIV | BD143114 | Red sea bream (*Pagrus major*) | Food fish, marine | Farmed | 1995 | Japan, Ehime Prefecture | 112,414 | 121 | Kurita et al., (2002); Nakajima et al. (1995); Zhang et al. (2013) |
|  |  |  | RBIV-C1 | KC244182 | Isolated from Spotted knifejaw (*Oplegnathus punctatus*)  Original strain isolated from farmed rock bream (*Oplegnathus fasciatus*) | Food fish, marine | Farmed fish | 2009 | China, Fujian Province | 112,333 | 118 | Zhang et al., (2014), Zhang et al. (2012) |
|  |  |  | GSIV-K1 | KT804738 | Barramundi (*Lates calcarifer*) | Food fish, marine or freshwater | Farmed | 2006 | Taiwan, Kaohsiung | 112,565 | 120 | Wen & Hong (2016) |
|  |  |  | PIV2016 | MK098187 | Pompano (*Trachinotus carolinus*) | Food fish, marine | Unknown | 2016 | Dominican Republic | 112,052 | 120 | Koda et al. (2019) |
|  |  |  | PIV2010 | MK098185 | Pompano (*Trachinotus carolinus*) | Food fish, marine | Unknown | 2010 | Dominican Republic | 112,321 | 120 | Koda et al. (2019) |
|  |  |  | PIV2014a | MK098186 | Pompano (*Trachinotus carolinus*) | Food fish, marine | Unknown | 2014 | Dominican Republic | 112,377 | 121 | Koda et al. (2019) |
|  |  |  | Zhoushan | MW139932 | Large yellow croaker (*Larimichthys crocea*) | Food fish, marine | Farmed, cage culture | 2020 | China, Zhoushan Island | 112,043 | 119 | Wang et al., (2022) |
|  |  |  | 17SbTy | OK042108 | Japanese seabass (*Lateolabrax japonicus)* | Food fish, marine | Farmed | 2017 | South Korea, Tongyeong | 112,360 | 121 | Jeong et al. (2021) |
|  |  |  | 17RbGs | OK042109 | Rock bream (*Oplegnathus fasciatus)* | Food fish, marine | Farmed | 2017 | South Korea, Goseong | 112,235 | 122 | Jeong et al. (2021) |
|  |  |  | SKIV-TJ | ON075463 | Spotted knifejaw (*Oplegnathus punctatus*) | Food fish, marine | Farmed | 2019 | China, Tianjin City | 112,489 | 120 | Liao et al., (2023) |
|  |  |  | SBIV-VP13 | ON740976 | Spotted seabass (*Lateolabrax maculatus*) | Food fish, marine | Farmed, pond culture | 2020 | China, Zhuhai, Guangdong Province | 112,158 | 120 | Fu et al. (2023) |
|  |  |  | SBIV-V12 | ON743042 | Spotted seabass (*Lateolabrax maculatus*) | Food fish, marine | Farmed, pond culture | 2020 | China, Zhuhai, Guangdong Province | 111,876 | 118 | Fu et al. (2023) |
|  |  |  | ZS-1 | OL774653 | large yellow croaker (*Larimichthys crocea)* | Food fish, marine | Unknown | 2020 | China, Zhoushan | 111,854 | 117 | GenBank |
|  |  |  | ZS-2 | OL774654 | large yellow croaker (*Larimichthys crocea*) | Food fish, marine | Unknown | 2020 | China, Zhoushan | 112,041 | 119 | GenBank |
|  |  |  | ZS-3 | OL774655 | large yellow croaker (*Larimichthys crocea*) | Food fish, marine | Unknown | 2020 | China,  Zhoushan | 112,362 | 119 | GenBank |
|  |  |  | RSIV_121 | MT798582 | Barramundi (*Lates calcarifer*) | Food fish, marine or freshwater | Farmed, open estuarine cages | 2018 | India, Mangalore | 111,557 | 116 | Puneeth et al., (2021), Girisha et al. (2020) |
|  |  |  | RSIV-HGIV-Cantik-2014 | MW883598 | Hybrid groper (*Epinephelus fuscoguttatus* x *Epinephelus polyphekadion*) | Food fish, marine | Unknown | 2014 | Indonesia, Bali | 112,717 | 120 | Fusianto et al., (2023) |
|  |  |  | RIE12-1 | AP017456 | Red sea bream (*Pagrus major*) | Food fish, marine | Unknown | 2012 | Japan,  Ehime prefecture | 112,590 | 121 | GenBank |
|  |  |  | Ehime-1 | AB104413 | Red sea bream (*Pagrus major*) | Food fish, marine | Unknown | 1990 | Japan | 112,415 | 121 | Nakajima and Kunita (2005), (Kurita et al. 2002) |
|  |  |  | FD201807 | OQ475017 | Large yellow croaker  (*Larimichthys crocea*) | Food fish, marine | Farmed | 2018 | China, Ningde, Fujian | 112,214 | 121 | Liu et al. (2025) |
|  |  | TRBIV | KEY15001 | MG570131 | Keyhole cichlid (*Cleithracara maronii*) | Ornamental fish, freshwater | Unknown | 1990 | United States | 111,347 | 118 | Kawato et al. (2021), Fusianto et al. (2023), Koda et al., (2018), Go et al., (2016) |
|  |  |  | TRT15003 | MG570132 | Three spotted gourami (*Trichopodus trichopterus*) | Ornamental fish, freshwater | Unknown | 1989 | United States | 111,591 | 119 | GenBank  Fusianto et al., (2023), Koda et al., (2018) |
|  |  |  | TRBIV | GQ273492 | Turbot (*Scophthalmus maximus*) | Food fish, marine | Farmed | 2005 | China, Shandong Province | 110,104 | 115 | Shi et al. (2010), Fusianto et al., (2023) |
|  | *Megalocytivirus lates1* | SDDV | SDDV-ZH-06/20 | OM037668 | Yellowfin seabream (*Acanthopagrus latus*) | Marine fish | Unknown | 2020 | China | 131,893 | 134 | Fu et al. (2025) |
|  |  |  | SDDV_Thai_2019 | MN562489 | Barramundi (*Lates calcarifer*) | Food fish, marine or freshwater | Unknown | 2018 | Thailand | 131,129 | 132 | GenBank |
|  |  |  | GF_MU1 | MT521409 | Barramundi (*Lates calcarifer*) | Food fish, marine or freshwater | Unknown | 2019 | Thailand | 131,276 | 132 | GenBank |
|  | Threespine stickleback iridovirus | N/A | TSIV | PQ335173_  PQ335174 | Threespine stickleback (*Gasterosteus aculeatus*) | Freshwater fish | Wild | 2012 | North America | 115,000 | 103 | Yoxsima et al. (2024) |
|  | European chub iridovirus | N/A | LEC15001 | MK637631 | European chub (*Squalius cephalus*) | Freshwater fish | Farmed | 2005 | United Kingdom | 128, 216 | 114 | Halaly et al. (2019) |
| *Ranavirus* | *Ranavirus rana 1* | N/A | FV3 | NC_005946 | Northern leopoard frog (*Lithobates pipiens*) | N/A | N/A | 1984 | Unknown | 105,903 | 97 | Granoff et al. (1965); Tan et al. (2004); Willis et al. (1984) |
|  | *Ranavirus epinephelus 1* | N/A | SGIV | AY521625 | Brown-spotted grouper (*Epinephelus chlorostigma*) | N/A | N/A | 1998 | Singapore | 140,131 | 141 | Song et al. (2004) |
| *Daphniairidovirus* | *Daphniairidovirus daphnia 1* | N/A | DIV-1 | LS484712 | Water flea (*Daphnia magna*) | N/A | N/A | 2017 | Finland | 288,858 | 278 | Toenshoff et al. (2018) |
| *Decapodiridovirus* | *Decapodiridovirus litopenaeus 1* | N/A | 20141215 | MF599468 | White leg shrimp (*Litopenaeus vannamei*) | N/A | N/A | 2014 | China | 165,809 | 174 | Qiu et al. (2017) |
| *Lymphocystivirus* | *Lymphocystivirus sparus 1* | N/A | SA9 | KX643370 | Gilthead sea bream  *(Sparus aurata*) | N/A | N/A | Between 2005 and 2015 | Spain | 208,501 | 152 | Lopez-Porras et al. (2018) |
|  | *Lymphocystivirus micropogonias 1* | N/A | LCDV-WC | MN803438 | Whitemouth croaker  (*Micropogonias furnieri*) | N/A | N/A | 2015 | Uruguay | 211,086 | 144 | Doszpoly et al. (2020) |
| *Iridovirus* | *Iridovirus chilo 1* | N/A | IIV-6 | AF303741 | Stem-boring lepidopteran (*Chilo suppressalis*) | N/A | N/A | 1987 | Germany | 212,482 | 203 | İnce et al., (2018); Jakob et al., (2001); Schnitzler et al. (1987) |
|  | *Iridovirus armadillidium 1* | N/A | IIV-31 | HF920637 | Pill bug, (*Armadillidium vulgare*) | N/A | N/A | 2010 | United States | 220,222 | 220 | Piégu et al. (2014) |
| *Chloriridovirus* | *Chloriridovirus simulium 1* | N/A | IIV-22 | HF920633 | Blackfly (*Simulium* spp.) | N/A | N/A | 1980 | Wales, United Kingdom | 204,815 | 179 | Piégu et al. (2014) |
|  | *Chloriridovirus anopheles 1* | N/A | AMIV | KF938901 | Mosquitoes  (*Anopheles minimus*) | N/A | N/A | 2007 | China | 163,023 | 157 | Huang et al., (2015) |
