## Supplementary table 2_Gene summary table for "Genomic analysis of *Megalocytivirus* genomes reveals widespread recombination"

**Supplementary table 2.** Summary of conserved genes reviewed in this study and comparison with other studies. Orthogroups marked with an asterisk were not included in the final trees as they contained insufficient genomic sequence information. The putative identity and orthogroup of putatively recombinant genes are bolded. Genome positions are based on the ISKNV genome (accession AF371960) reannotated in this study available at <https://github.com/PollyHannah/Phylogenomic-study/blob/main/annotated_genome_sequences.geneious>. This is except for gene number 20 which is based on AFIV-16 genome (accession MK689685) re-annotated in this study and available via the same link. **N°** Number **RE** Recombination evidence **MCV BL** Megalocytivirus genus branch length **-** not applicable.

|  |  |  | |  | |  |  | |  | |  | **Previous studies** | | |
| --- | --- | --- | --- | --- | --- | --- | --- | --- | --- | --- | --- | --- | --- | --- |
| **N°** | **Putative identity** | **Position**^[1]^ | | **Family** | | | **Genus** | | **Species** | |  | **Eaton et al. (2007)** | **Zhao et al. (2023)** | **Fusianto et al. (2023)** |
|  |  | **Start** | **Stop** | **Orthogroup** | **RE** | **MCV BL** | **Orthogroup** | **RE** | **Orthogroup** | **RE** | **Intragenic or**  **intergenic**  **recombination** |  |  |  |
| 1 | **Sodium-coupled neutral amino acid symporter 2** | 134 | 1270 | OG0000030* | - | - | OG0000049 | No | **OG0000084** | **Yes** | Intragenic | - | - | 1 |
| 2 | DNA-directed RNA polymerase subunit | 1240 | 1695 | OG0000031* | - | - | OG0000001 | No | OG0000002 | No | - | - | - | 2 |
| 3 | Caspase-1 | 1772 | 2077 | OG0000086* | - | - | OG0000086* | - | OG0000003 | No | - | - | - | - |
| 4 | B2 bradykinin receptor | 2077 | 2568 | OG0000101* | - | - | OG0000101* | - | OG0000085 | No | - | - | - | - |
| 5 | NIF NLI interacting factor | 2893 | 3453 | OG0000004 | No | 0.70 | OG0000002 | No | OG0000004 | No | - | 5L | Cg8 | - |
| 6 | E3 ubiquitin-protein ligase | 3485 | 3790 | OG0000028* | - | - | OG0000050 | No | OG0000086 | No | - | - | - | - |
| 7 | Major capsid protein | 3794 | 5155 | OG0000005 | No | 0.40 | OG0000003 | No | OG0000005 | No | - | 6L | Cg16 | 7 |
| 8 | **Myristoylated protein** | 5174 | 6631 | OG0000001 | No | 0.52 | OG0000004 | No | **OG0000006** | **Yes** | Intergenic | 7L | Cg10 | 8 |
| 9 | **Auxin transport protein** | 6702 | 8246 | OG0000065* | - | - | OG0000063 | No | **OG0000007** | **Yes** | Intergenic | - | - | - |
| 10 | **CCA-adding enzyme** | 8200 | 8418 | OG0000061* | - | - | OG0000061 | No | **OG0000104** | **Yes** | Intragenic | - | - | - |
| 11 | **DNA polymerase alpha catalytic subunit** | 8662 | 9054 | OG0000066* | - | - | OG0000064 | No | **OG0000008** | **Yes** | Intragenic | - | - | 11 |
| 12 | **Carboxy-S-adenosyl-L-methionine synthase** | 9051 | 9311 | OG0000053* | - | - | OG0000051 | No | **OG0000009** | **Yes** | Intergenic | - | - | 12 |
| 13 | **E3 ubiquitin-protein ligase** | 9330 | 9662 | OG0000032* | - | - | OG0000005 | No | **OG0000010** | **Yes** | Intragenic | - | - | 13 |
| 14 | **Serine/threonine-protein kinase** | 9669 | 11054 | OG0000002 | No | - | OG0000000 | No | **OG0000011** | **Yes** | Intragenic | 13R | - | - |
| 15 | Nucleoside triphosphate pyrophosphatase | 11017 | 11310 | - | - | - | OG0000115 | No | OG0000115 | No | - | - | - | ORF15 |
| 16 | **Ketol-acid reductoisomerase (NADP(+))** | 11309 | 12268 | OG0000067* | - | - | OG0000065 | No | **OG0000012** | **Yes** | Intragenic | - | - | 15 |
| 17 | **Bifunctional protein FolD** | 12278 | 13069 | OG0000033* | - | - | OG0000006 | No | **OG0000013** | **Yes** | Intergenic | - | - | 16 |
| 18 | **tRNA (cytidine(56)-2'-O)-methyltransferase** | 13129 | 13716 | OG0000034* | - | - | OG0000007 | No | **OG0000014** | **Yes** | Intergenic | - | - | 17 |
| 19 | **Uncharacterised protein** | 13731 | 14063 | OG0000035* | - | - | OG0000008 | No | **OG0000015** | **Yes** | Intergenic | - | - | - |
| 20 | **S-adenosylmethionine:tRNA ribosyltransferase-isomerase** | 14089 | 14325 | OG0000113* | - | - | OG0000113 | No | **OG0000112** | **Yes** | Intergenic | - | - | No match |
| 21 | **Neurofilament heavy polypeptide** | 14322 | 14513 | OG0000085* | - | - | OG0000085 | No | **OG0000106** | **Yes** | Intergenic | - | - | - |
| 22 | DNA polymerase | 14579 | 17425 | OG0000006 | No | 0.78 | OG0000009 | No | OG0000016 | No | - | 19R | Cg11 | 20 |
| 23 | Aspartate carbamoyltransferase catalytic subunit | 17467 | 18006 | OG0000036* | - | - | OG0000010 | No | OG0000017 | No | - | - | - | 21 |
| 24 | **ADP-ribose glycohydrolase** | 17990 | 19489 | OG0000054* | - | - | OG0000052 | No | **OG0000087** | **Yes** | Intergenic | - | - | - |
| 25 | **Ribonucleoside-diphosphate reductase small subunit** | 22300 | 23238 | **OG0000017** | **Yes** | 2.67 | OG0000011 | No | **OG0000018** | **Yes** | Intragenic | 24R | - | 24 |
| 26 | **UPF0319 protein HSM_0266** | 23822 | 24145 | OG0000102* | - | - | OG0000102* | - | **OG0000088** | **Yes** | Intragenic | - | - | - |
| 27 | **DNA repair protein RAD2** | 24167 | 25063 | OG0000003 | No | 0.61 | OG0000012 | No | **OG0000019** | **Yes** | Intragenic | 28L[1] | Cg17 | 27 |
| 28 | **DNA-directed RNA polymerase** | 25080 | 28586 | OG0000000 | No | 0.62 | OG0000013 | No | **OG0000020** | **Yes** | Intragenic | 28L | Cg2 | 28 |
| 29 | **Transcription factor S** | 28593 | 28814 | **OG0000007** | **Yes** | 1.81 | OG0000014 | No | **OG0000021** | **Yes** | Intragenic | 29L | - | 29 |
| 30 | **Agmatinylcytidine synthetase TiaS** | 28884 | 29486 | OG0000062* | - | - | OG0000062 | No | **OG0000089** | **Yes** | Intragenic | - | - | - |
| 31 | **Kinase protein** | 29447 | 30061 | OG0000008 | No | 1.00 | OG0000015 | No | **OG0000022** | **Yes** | Intragenic | 34R[2] | Newcg4 | - |
| 32 | **2-(5''-triphosphoribosyl)-3'-dephosphocoenzyme-A synthase** | 30138 | 31040 | OG0000103* | - | - | OG0000103* | - | **OG0000090** | **Yes** | Intragenic | - | - | - |
| 33 | **DNA-directed RNA polymerase II subunit** | 31144 | 34278 | OG0000009 | No | 0.57 | OG0000053 | No | **OG0000091** | **Yes** | Intergenic | 34R | Cg12 | 33 |
| 34 | **ATP-dependent helicase/nuclease subunit A** | 34360 | 35493 | OG0000055* | - | - | OG0000054 | No | **OG0000023** | **Yes** | Intragenic | - | - | - |
| 35 | **Uncharacterised protein** | 35576 | 36601 | OG0000087* | - | - | OG0000087 | No | **OG0000092** | **Yes** | Intragenic | - | - | - |
| 36 | Uncharacterised protein | 36598 | 37950 | OG0000106* | - | - | OG0000106 | No | OG0000107 | No | - | - | - | - |
| 37 | **Glutamyl-tRNA(Gln) amidotransferase subunit F** | 37959 | 39395 | OG0000104* | - | - | OG0000104* | - | **OG0000093** | **Yes** | Intragenic | - | - | - |
| 38 | Uncharacterised protein | 39439 | 40311 | OG0000068* | - | - | OG0000066 | No | OG0000024 | No | - | - | - | - |
| 39 | Uncharacterised protein | 40304 | 41443 | OG0000069* | - | - | OG0000067 | No | OG0000025 | No | - | - | - | - |
| 40 | **Probable succinyl-CoA:3-ketoacid coenzyme A transferase** | 41445 | 42788 | OG0000088* | - | - | OG0000088 | No | **OG0000094** | **Yes** | Intergenic | - | - | - |
| 41 | **Uncharacterised protein** | 42803 | 43396 | OG0000070* | - | - | OG0000068 | No | **OG0000026** | **Yes** | Intergenic | - | - | - |
| 42 | **Erv1 / Alr family** | 43480 | 43842 | OG0000022 | No | 0.50 | OG0000016 | No | **OG0000027** | **Yes** | Intergenic | 43L | Newcg5 | 42 |
| 43 | **Protein translocase subunit SecD** | 43845 | 44645 | OG0000071* | - | - | OG0000069 | No | **OG0000028** | **Yes** | Intergenic | - | - | - |
| 44 | **Uncharacterised protein** | 44650 | 45564 | OG0000072* | - | - | OG0000070 | No | **OG0000029** | **Yes** | Intergenic | - | - | - |
| 45 | **Probable DNA (cytosine-5)-methyltransferase** | 45558 | 46241 | OG0000027 | No | 1.03 | OG0000017 | No | **OG0000030** | **Yes** | Intergenic | - | - | 45 |
| 46 | **Uncharacterised protein** | 46401 | 46664 | OG0000037* | - | - | OG0000018 | No | **OG0000031** | **Yes** | Intergenic | - | - | 46 |
| 47 | **Vascular endothelial growth factor** | 46661 | 47005 | OG0000038* | - | - | OG0000019 | No | **OG0000032** | **Yes** | Intergenic | - | - | - |
| 48 | **Folylpolyglutamate synthase** | 47021 | 47191 | OG0000073* | - | - | OG0000071 | No | **OG0000033** | **Yes** | Intergenic | - | - | 48 |
| 49 | **Transcription factor SFL2** | 47250 | 47678 | OG0000074* | - | - | OG0000072 | No | **OG0000034** | **Yes** | Intergenic | - | - | 49 |
| 50 | **Uncharacterised protein** | 47951 | 48403 | OG0000089* | - | - | OG0000089* | - | **OG0000035** | **Yes** | Intergenic | - | - | - |
| 51 | **Cytosol aminopeptidase** | 48633 | 49559 | OG0000039* | - | - | OG0000020 | No | **OG0000036** | **Yes** | Intergenic | - | - | - |
| 52 | **Serine/threonine-protein kinase** | 49582 | 50508 | OG0000040* | - | - | OG0000021 | No | **OG0000037** | **Yes** | Intergenic | 55L | - | - |
| 53 | **Uncharacterised protein** | 50519 | 51166 | OG0000010 | No | 0.68 | OG0000022 | No | **OG0000038** | **Yes** | Intergenic | 56L | Cg5 | 54 |
| 54 | **Electron transfer flavoprotein** | 51173 | 51433 | OG0000029* | - | - | OG0000023 | No | **OG0000039** | **Yes** | Intergenic | - | - | 55 |
| 55 | **4-hydroxy-tetrahydrodipicolinate reductase** | 51749 | 52414 | OG0000114* | - | - | OG0000114 | No | **OG0000114** | **Yes** | Intergenic | - | - | No match |
| 56 | **Replication factor** | 52359 | 53162 | OG0000041* | - | - | OG0000024 | No | **OG0000040** | **Yes** | Intergenic | 61L | Cg1 | 58 |
| 57 | **Titin homolog** | 53159 | 56785 | OG0000109* | - | - | OG0000109 | No | **OG0000113** | **Yes** | Intergenic | - | - | - |
| 58 | **Transaminase htyB** | 56845 | 57213 | OG0000075* | - | - | OG0000073 | No | **OG0000041** | **Yes** | Intragenic | - | - | 60 |
| 59 | **Helicase protein** | 57227 | 59875 | OG0000011 | No | 0.77 | OG0000025 | No | **OG0000042** | **Yes** | Intragenic | 63L | Cg3 | 61 |
| 60 | **mRNA-capping enzyme** | 59918 | 61393 | OG0000024* | - | - | OG0000026 | No | **OG0000043** | **Yes** | Intergenic | - | - | 62 |
| 61 | **E3 ubiquitin-protein ligase** | 61439 | 61900 | OG0000084* | - | - | OG0000084* | - | **OG0000001** | **Yes** | Intergenic | - | - | - |
| 62 | **RING finger protein** | 61982 | 63025 | OG0000064* | - | - | OG0000083 | No | **OG0000082** | **Yes** | Intergenic | - | - | - |
| 63 | **D83 antigen** | 63271 | 63855 | OG0000076* | - | - | OG0000074 | No | **OG0000044** | **Yes** | Intergenic | - | - | - |
| 64 | **Uncharacterised protein** | 63896 | 65329 | OG0000090* | - | - | OG0000090 | No | **OG0000095** | **Yes** | Intergenic | - | - | 66 |
| 65 | **tRNA-specific 2-thiouridylase MnmA** | 65336 | 66001 | OG0000107* | - | - | OG0000107* | - | **OG0000105** | **Yes** | Intergenic | - | - | - |
| 66 | **Urotensin** | 66062 | 66538 | OG0000110* | - | - | OG0000110* | - | **OG0000108** | **Yes** | Intergenic | - | - | - |
| 67 | **Demethylmenaquinone methyltransferase** | 66432 | 68042 | OG0000091* | - | - | OG0000091 | No | **OG0000096** | **Yes** | Intergenic | - | - | 69 |
| 68 | **Genome polyprotein** | 69203 | 69622 | OG0000092* | - | - | OG0000092* | - | **OG0000045** | **Yes** | Intergenic | - | - | - |
| 69 | **Lissencephaly-1 homolog** | 69669 | 70682 | OG0000093* | - | - | OG0000093* | - | **OG0000046** | **Yes** | Intergenic | - | - | - |
| 70 | Recombination protein RecR | 70777 | 71043 | OG0000112* | - | - | OG0000112* | - | OG0000110 | No | - | - | - | - |
| 71 | **Uncharacterised protein** | 71045 | 74017 | OG0000018 | No | 0.57 | OG0000055 | No | **OG0000097** | **Yes** | Intragenic | 76L | Cg9 | 74 |
| 72 | **Ankyrin repeat protein** | 74035 | 75369 | OG0000077* | - | - | OG0000075 | No | **OG0000047** | **Yes** | Intragenic | - | - | 75 |
| 73 | **Branched-chain-amino-acid aminotransferase** | 75366 | 75830 | OG0000042* | - | - | OG0000027 | No | **OG0000048** | **Yes** | Intragenic | - | - | 76 |
| 74 | **Homeodomain-interacting protein kinase 2** | 75832 | 76053 | OG0000094* | - | - | OG0000094* | - | **OG0000049** | **Yes** | Intragenic | - | - | - |
| 75 | **Uncharacterised protein** | 76367 | 76864 | OG0000043* | - | - | OG0000028 | No | **OG0000050** | **Yes** | Intergenic | - | - | 78 |
| 76 | **mRNA-capping enzyme** | 76901 | 78007 | OG0000044* | - | - | OG0000029 | No | **OG0000051** | **Yes** | Intragenic | - | - | 79 |
| 77 | **Uncharacterised protein** | 78059 | 78418 | OG0000045* | - | - | OG0000030 | No | **OG0000052** | **Yes** | Intergenic | - | - | 80 |
| 78 | **RNA replication polyprotein** | 78526 | 79881 | OG0000078* | - | - | OG0000076 | No | **OG0000053** | **Yes** | Intergenic | - | - | - |
| 79 | **ATP-dependent RNA helicase** | 79884 | 80486 | OG0000079* | - | - | OG0000077 | No | **OG0000054** | **Yes** | Intergenic | 82R[4] | - | - |
| 80 | **Uncharacterised protein** | 80483 | 80947 | OG0000012 | No | 0.38 | OG0000031 | No | **OG0000055** | **Yes** | Intergenic | - | Newcg7 | - |
| 81 | **Ribonuclease 3** | 80940 | 81710 | OG0000019 | No | 1.38 | OG0000032 | No | **OG0000056** | **Yes** | Intragenic | 87R | - | 84 |
| 82 | **Putative SAP domain-containing protein** | 81707 | 82120 | OG0000025 | No | 0.76 | OG0000033 | No | **OG0000057** | **Yes** | Intragenic | - | - | 85 |
| 83 | **Uncharacterised protein** | 82164 | 83720 | OG0000046* | - | - | OG0000034 | No | **OG0000058** | **Yes** | Intergenic | - | - | ORF88 |
| 84 | **Membrane protein** | 83701 | 84663 | OG0000013 | No | 0.35 | OG0000035 | No | **OG0000059** | **Yes** | Intragenic | 85L[3] | - | - |
| 85 | **Cell division protein ZipA** | 84702 | 84863 | OG0000056* | - | - | OG0000056 | No | **OG0000098** | **Yes** | Intergenic | - | - | - |
| 86 | **Zinc finger CCCH-type antiviral protein 1** | 84860 | 85786 | OG0000057* | - | - | OG0000057 | No | **OG0000099** | **Yes** | Intergenic | - | - | 91 |
| 87 | **Thiamine-phosphate synthase** | 85796 | 86296 | OG0000083* | - | - | OG0000082* | - | **OG0000000** | **Yes** | Intergenic | - | - | 92 |
| 88 | **Uncharacterised protein** | 86321 | 87481 | OG0000047* | - | - | OG0000036 | No | **OG0000060** | **Yes** | Intragenic | - | - | - |
| 89 | **Uncharacterised protein** | 87489 | 88226 | OG0000020 | No | 0.58 | OG0000037 | No | **OG0000061** | **Yes** | Intragenic | 96L | Newcg2 | 94 |
| 90 | **Uncharacterised protein** | 88232 | 88687 | OG0000048* | - | - | OG0000038 | No | **OG0000062** | **Yes** | Intragenic | - | - | 95 |
| 91 | **Uncharacterised protein** | 88774 | 89097 | OG0000058* | - | - | OG0000058 | No | **OG0000063** | **Yes** | Intragenic | - | - | - |
| 92 | **Uncharacterised protein** | 89144 | 89689 | OG0000063* | - | - | OG0000078 | No | **OG0000064** | **Yes** | Intragenic | - | - | - |
| 93 | **Auxin transport protein BIG** | 89736 | 90251 | OG0000049* | - | - | OG0000039 | No | **OG0000065** | **Yes** | Intragenic | - | - | 98 |
| 94 | **E3 ubiquitin-protein ligase HACE1** | 90311 | 91753 | OG0000080* | - | - | OG0000079 | No | **OG0000066** | **Yes** | Intragenic | - | - | - |
| 95 | **Suppressor of cytokine signaling 1** | 91760 | 92161 | OG0000095* | - | - | OG0000095* | - | **OG0000067** | **Yes** | Intragenic | - | - | - |
| 96 | **Uncharacterised protein** | 92215 | 92991 | OG0000081* | - | - | OG0000080 | No | **OG0000068** | **Yes** | Intragenic | - | - | 101 |
| 97 | **Succinyl-5-enolpyruvyl-6-hydroxy-3-cyclohexene-1-carboxylate** | 92993 | 93358 | OG0000096* | - | - | OG0000096* | - | **OG0000069** | **Yes** | Intragenic | - | - | 102 |
| 98 | **Replication protein E1** | 93482 | 94501 | OG0000100* | - | - | OG0000100* | - | **OG0000083** | **Yes** | Intragenic | - | - | - |
| 99 | **Sodium/potassium-transporting ATPase subunit alpha-4** | 94494 | 95093 | OG0000105* | - | - | OG0000105* | - | **OG0000100** | **Yes** | Intergenic | - | - | - |
| 100 | **D5 family NTPase** | 95185 | 97950 | OG0000014 | No | 0.58 | OG0000040 | No | **OG0000070** | **Yes** | Intragenic | 109L | Cg6 | 105 |
| 101 | **Lipopolysaccharide core galacturonosyltransferase** | 97997 | 98152 | OG0000097* | - | - | OG0000097* | - | **OG0000071** | **Yes** | Intragenic | - | - | 106 |
| 102 | **TNF receptor-associated factor 2** | 98149 | 99039 | OG0000098* | - | - | OG0000098 | No | **OG0000101** | **Yes** | Intragenic | - | - | - |
| 103 | **Proliferating cell nuclear antigen** | 99059 | 99802 | OG0000026 | No | 0.2 | OG0000041 | No | **OG0000072** | **Yes** | Intragenic | 112R | - | 108 |
| 104 | Polyketide synthase 14 | 99792 | 100295 | OG0000050* | - | - | OG0000042 | No | OG0000073 | No | - | - | - | 109 |
| 105 | **Kinase** | 100334 | 102916 | **OG0000021** | **Yes** | 1.67 | OG0000043 | No | **OG0000074** | **Yes** | Intragenic | 114L | - | - |
| 106 | **Immediate-early protein ICP-46** | 103203 | 104213 | OG0000023 | No | 0.61 | OG0000044 | No | **OG0000075** | **Yes** | Intragenic | 115R | Newcg6 | 111 |
| 107 | **Microtubule-actin cross-linking factor 1** | 104276 | 105667 | OG0000111* | - | - | OG0000111 | No | **OG0000111** | **Yes** | Intragenic | - | - | - |
| 108 | **Early 31 kDa protein** | 105721 | 106335 | OG0000051* | - | - | OG0000045 | No | **OG0000076** | **Yes** | Intragenic | - | - | 114 |
| 109 | **Vacuolar membrane protease** | 106492 | 106752 | OG0000099* | - | - | OG0000099* | - | **OG0000077** | **Yes** | Intragenic | - | - | - |
| 110 | **Ankyrin repeat protein** | 106723 | 108042 | OG0000059* | - | - | OG0000059 | No | **OG0000078** | **Yes** | Intragenic | - | - | - |
| 111 | **E3 ubiquitin-protein ligase** | 108105 | 108392 | OG0000082* | - | - | OG0000081 | No | **OG0000102** | **Yes** | Intragenic | - | - | 116 |
| 112 | **Uncharacterised protein** | 108424 | 108933 | OG0000052* | - | - | OG0000046 | No | **OG0000079** | **Yes** | Intragenic | - | - | 117 |
| 113 | **Uncharacterised protein** | 108934 | 109584 | OG0000108* | - | - | OG0000108 | No | **OG0000109** | **Yes** | Intragenic | - | - | - |
| 114 | **ATPase** | 109594 | 110313 | OG0000015 | No | 0.36 | OG0000047 | No | **OG0000080** | **Yes** | Intragenic | 122R | Cg4 | 119 |
| 115 | **Uncharacterised protein** | 110285 | 110656 | OG0000016 | No | 0.48 | OG0000048 | No | **OG0000081** | **Yes** | Intragenic | - | - | 120 |
| 116 | **ATP-dependent DNA helicase RecG** | 110665 | 111351 | OG0000060* | - | - | OG0000060 | No | **OG0000103** | **Yes** | Intragenic | - | - | 121 |
| **Total number of genes retained** | | | | **24** | | | **96** | | **68** | |  |  |  |  |
