## Supplementary Materials for "Genomic analysis of *Megalocytivirus* genomes reveals widespread recombination"

### RDP4 recombination detection

A conservative set of four criteria (as outlined in the main manuscript) were used to refine each recombination hypothesis generated by RDP4 (version 101) (Martin et al., 2015) for *Megalocytivirus pagrus1*. Example plots are provided below, which were used to accept a recombination event within the infectious spleen and kidney necrosis (ISKNV) genome RSIV-Ku (accession KT781098) (Figure 1, Figure 2, Figure 3).

#### RDP4 MAXCHI and CHIMAREA plots

For a recombination event generated by RDP4 (version 101) (Martin et al., 2015) to be accepted, the recombination signal needed to be visible in the MAXCHI (Smith, 1992) (Figure 1) and CHIMAREA (Posada & Crandall, 2001) (Figure 2) plots generated by RDP4 (version 101) (Martin et al., 2015).

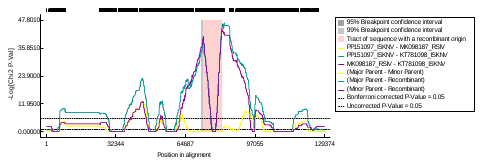

Figure 1. MAXCHI plot generated by RDP4 (version 101) for the putative recombination event between infectious spleen and kidney necrosis virus genome RSIV-Ku (accession KT781098) and a red sea bream iridovirus (RSIV) clade 1 genome. The borders of the potential recombinant region (highlighted in red), as identified by GENECONV in RDP4, match up with the peaks of the purple and green lines, confirming the recombination signal and breakpoint positions for a putative recombination event between RSIV-Ku genome (accession KT781098) and RSIV clade 1 genome PIV2016 (accession MK098187) at a specified region in the whole genome alignment.

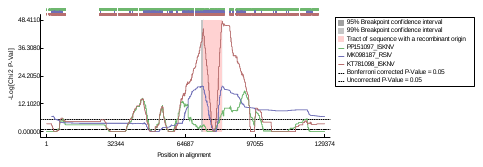

Figure 2. CHIMAREA plot generated by RDP4 (version 101) showing a recombination signal and breakpoint positions for recombination within the ISKNV genome RSIV-Ku (accession KT781098). The borders of the putative recombinant region (highlighted in red), as identified by GENECONV in RDP4, match up with the peak of the red line, indicating the existence of a recombination event within the RSIV-Ku genome (accession KT781098) at a specified region in the whole genome alignment.

#### RDP4 phylogenetic trees

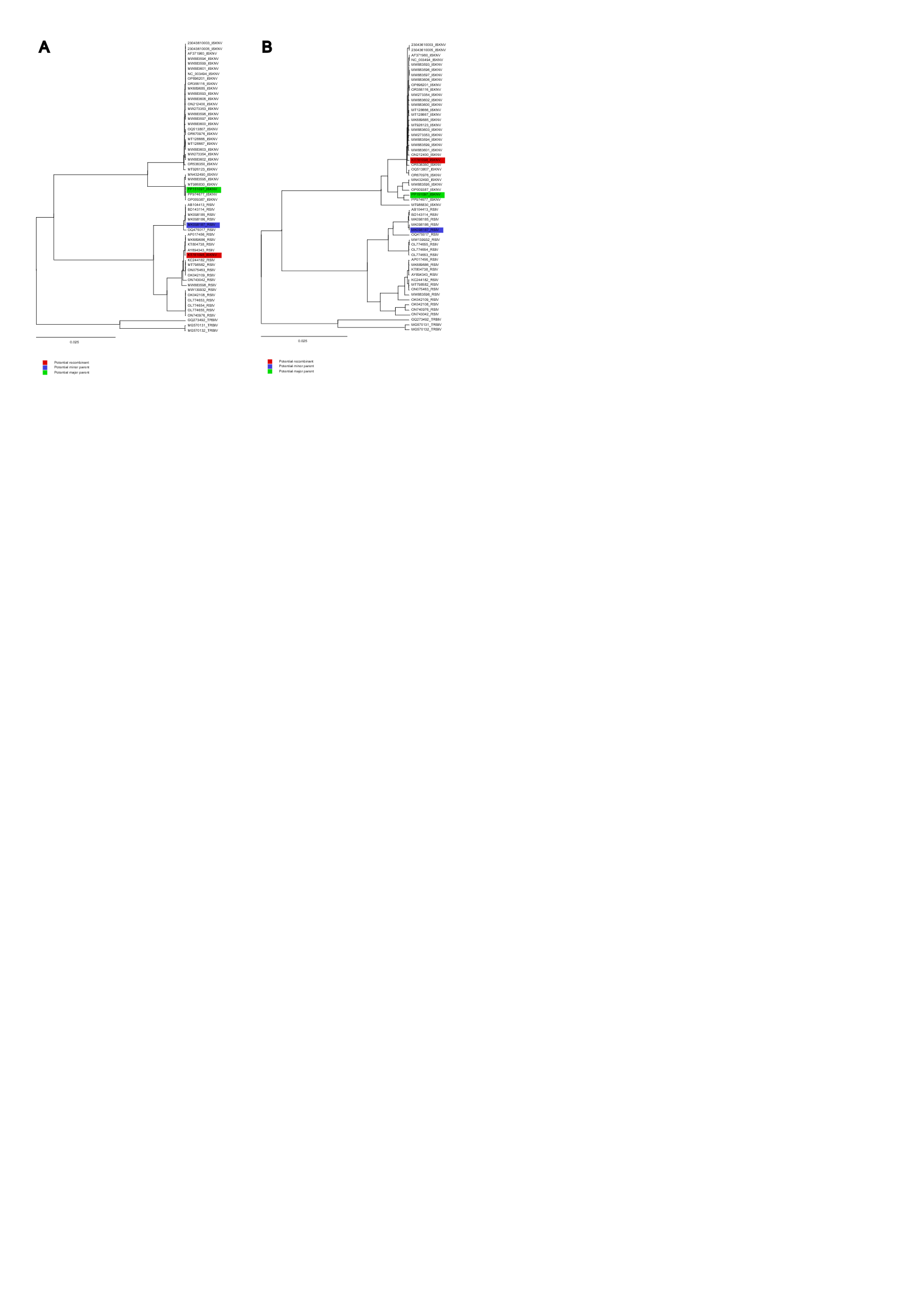
For a recombination event generated by RDP4 (version 101) (Martin et al., 2015) to be accepted, the recombination signal needed to be observed by comparing the topology of two phylogenetic trees generated by RDP4 – one based on the putative recombinant region and the other on combined regions where recombination was not detected (Figure 3).

Figure 3. Phylogenetic trees generated by generated by RDP4 (version 101) for a putative recombination event between infectious spleen and kidney necrosis virus (ISKNV) genome RSIV-Ku (accession KT781098) and a red sea bream iridovirus (RSIV) genome. The scale bars indicate the number of amino acid substitutions per site. A) phylogenetic tree including only the putative recombinant region from the whole genome alignment. B) Phylogenetic tree including only the concatenated regions outside of the putative recombinant region within the whole genome alignment. The topology of the trees A and B are different, indicating recombination. The putative recombinant ISKNV genome RSIV-Ku (accession KT781098) clusters with RSIV genomes in tree A, compared to ISKNV in tree B, indicating recombination between ISKNV genome RSIV-Ku (accession KT781098) and RSIV.

### Recombination analysis: Family trees

Four genes showed evidence of recombination at the family-level. Phylogenetic analysis of the putative serine/threonine-protein kinase gene revealed paraphyly consistent with the possible mixing of genetic information between members of different genera part of the *Iridoviridae* family (Figure 4). An AU test showed that we cannot reject monophyly and there is no evidence of recombination at this locus. Further detail on this analysis can be found on GitHub at this link: [Phylogenomic-study/iqtree_AU_test at main · PollyHannah/Phylogenomic-study](https://github.com/PollyHannah/Phylogenomic-study/tree/main/iqtree_AU_test).

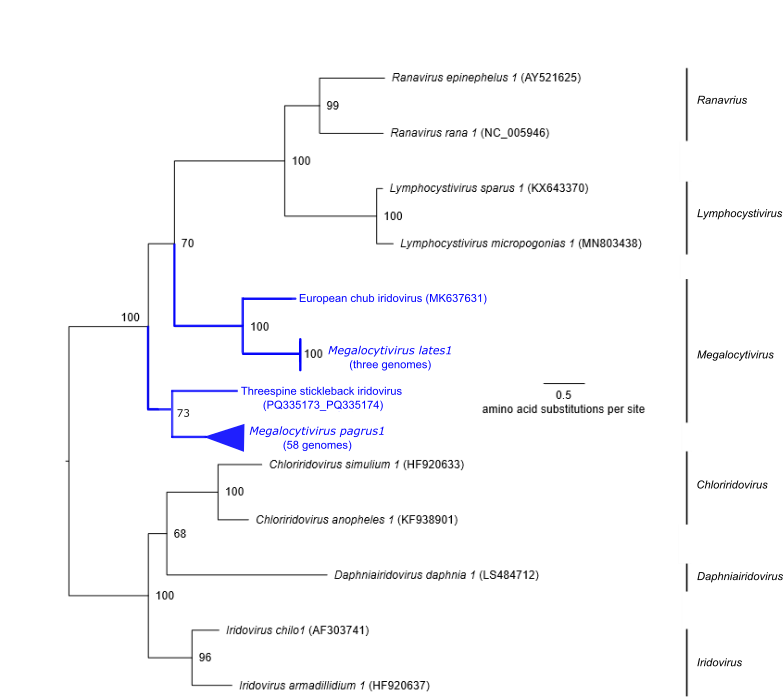

Figure 4. Phylogenetic tree of the putative serine/threonine-protein kinase gene. Maximum

likelihood analysis based on 73 genomes part of the family *Iridoviridae*. Bootstrap values are provided. The bootstrap value of the node which causes paraphyly of the megalocytiviruses is 70%. An AU test confirms that forcing the megalocytiviruses to be monophyletic produces a tree that is not significantly different from the maximum likelihood tree shown in this figure. This means that we cannot reject the hypothesis that the paraphyly is due to tree reconstruction uncertainty.

Consistent with Kurita and Nakajima (2012), analysis of the putative *Megalocytivirus* ribonucleoside-diphosphate reductase small subunit gene (RR-2) shows it is more closely related to eukaryotic organisms (including fish *Sardina pilchardus, Alosa alosa* and *Limanda limanda*, an annelid worm *Capitella teleta* and clam *Spisula solidissima*), than the remaining members of the family *Iridoviridae* (supplementary data_RR-2 gene_BLAST+.csv). This indicates that megalocytiviruses acquired their RR-2 gene from a past eukaryotic host (Kurita and Nakajima, 2012). The family-level RR-2 gene phylogeny reflects this divergence, with *Megalocytivirus* taxa positioned on an unusually long branch length of 2.67 amino acid substitutions per site (supplementary table 2).

Similarly, the putative transcription factor S gene in megalocytiviruses sits on a long branch of 1.81 amino acid substitutions per site (supplementary table 2), and the BLAST+ search revealed it shares a greater level of identity to eukaryotic organisms such as pipefish *Syngnathus scovelli,* protists (*Blastocystis spp.*), and fungi (*Rhizophagus spp*.), compared to that of other *Iridoviridae* genomes (supplementary data_Transcription factor S_BLAST+.csv).

A BLAST+ search of the putative kinase gene in megalocytiviruses indicates that it is more closely related to bacteria (including *Celeribacter spp.* and *Paracoccus zhejiangensis*) than other members of the *Iridoviridae* (supplementary data_Kinase gene_BLAST+.csv). This is consistent with the swapping of genetic information between megalocytiviruses and divergent organisms. Unsurprisingly, phylogenetic analysis of the putative kinase gene shows megalocytiviruses to be evolutionary divergent to all other *Iridoviridae* taxa, sitting on a long branch of 1.67 amino acid substitutions per site (supplementary table 2).

All family-level gene trees generated as part of this study can be found on GitHub at this link: <https://github.com/PollyHannah/Phylogenomic-study/tree/main/iqtree_family_trees>. The putative identify and orthogroup of each gene can be found in supplementary table 2.
